# *In vitro* infection of human lung tissue with SARS-CoV-2: Heterogeneity in host defense and therapeutic response

**DOI:** 10.1101/2021.01.20.427541

**Authors:** Matthew A. Schaller, Yamini Sharma, Zadia Dupee, Duy Nguyen, Juan Uruena, Ryan Smolchek, Julia C. Loeb, Tiago N. Machuca, John A. Lednicky, David J. Odde, Robert F. Campbell, W. Gregory Sawyer, Borna Mehrad

**Affiliations:** Division of Pulmonary, Critical Care, and Sleep Medicine, Gainesville, FL; Department of Mechanical and Aerospace Engineering, Gainesville, FL; Department of Environmental and Global Health, and Emerging Pathogens Institute, Gainesville, FL; Division of Cardiothoracic Surgery, University of Florida, Gainesville, FL; Department of Biomedical Engineering, University of Minnesota, Minneapolis, MN; Department of Drug Development, Walter Reed Army Institute of Research, Silver Spring, MD

## Abstract

Cell lines are the mainstay in understanding the biology of COVID-19 infection, but do not recapitulate many of the complexities of human infection. The use of human lung tissue is one solution for the study of such novel respiratory pathogens. We hypothesized that a cryopreserved bank of human lung tissue allows for the *in vitro* study of the inter-individual heterogeneity of host response to SARS-CoV-2 infection, thus providing a bridge between studies with cell lines and studies in animal models. We generated a cryobank of tissues from 16 donors, most of whom had risk factors for severe illness from COVID-19. Cryopreserved tissues preserved 90% of cell viability and contained heterogeneous populations of metabolically active epithelial, endothelial, and immune cell subsets of the human lung. Samples were readily infectible with HCoV-OC43 and SARS-CoV-2 coronavirus strains, and demonstrated comparable susceptibility to infection. In contrast, we observed a marked donor-dependent heterogeneity in the expression of IL-6, CXCL8 and IFNβ in response to SARS-CoV-2 infection. Treatment of tissues with dexamethasone and the experimental drug, N-hydroxycytidine, suppressed viral growth in all samples, whereas chloroquine and remdesivir had no detectable effect. Metformin and sirolimus, molecules with predicted antiviral activity, suppressed viral replication in tissues from a subset of donors. In summary, we developed a novel system for the *in vitro* study of human SARS-CoV-2 infection using primary human lung tissue from a library of donor tissues. This model may be useful for drug screening and for understanding basic mechanisms of COVID-19 pathogenesis.

**Importance:** The current biological systems for the study of COVID-19 are *in vitro* systems that differ from the human lung in many respects, and animal hosts to which the virus is not adapted. We developed another alternative for studying pathogenesis and drug susceptibility of SARS-CoV-2 in a cryopreserved bank of human lung tissues. We consider the importance of this work to relate to the practical use of this culture system as a repeatable and scalable approach that allows for the study of an important infection in relevant tissues.

The tissue bank highlights the heterogeneous response to SARS-CoV-2 infection and treatment, which allows researchers to investigate why treatments work in some donors but not others.

## Introduction

The COVID-19 pandemic has highlighted the need for improved models of infection to study host-pathogen interactions, rapidly screen potential therapeutic interventions, and study fundamental pathogenic mechanisms. Cell lines have served as the mainstay in understanding the biology of the infection, but do not capture the cellular heterogeneity, cell-cell and cell-matrix interactions, or between-host variability encountered in human infection.

The use of human lung tissue is one solution for the study of biology of novel respiratory pathogens. Lung tissue explants capture the cellular heterogeneity within the human lung and the use of tissues from multiple donors can allow the study of the genetic diversity found within the human population. To this end, human lung organoids (1, 2) and lung-on-a-chip organs (3) are readily infected with SARS-CoV-2. The use of these models is limited by two considerations: first, there are practical difficulties of repeating experiments with the same donor or performing experiments on multiple donors in parallel. Second, these are simplified systems that do not recapitulate the intrinsic complexities of lung tissue. The optimal solution to these problems is a repeatable and scalable *in vitro* system that captures the intricacies of the human lung while allowing high-throughput parallel drug screens from multiple donors and sequential experiments using tissue from the same donor.

Polyampholytes are polymeric hydrogels composed of poly-L lysine or other amphiphilic chemicals that can serve as a less toxic alternative to dimethylsulfoxide for the cryopreservation of cells. Similar to antifreeze proteins first discovered in arctic fish (4), these amphiphilic proteins interact with both the hydrophobic cell membrane and hydrophilic ice crystals, mitigating damage associated with the formation of crystals during freezing (5–7). Polyampholyte medias have been used to cryopreserve many cell lines and primary cells with high viability (8). Recent work has demonstrated that they can also be used to cryopreserve 3D cell structures (9). We therefore tested the hypothesis that a cryopreserved bank of human lung tissue allows for the *in vitro* study of the inter-individual heterogeneity of host response to SARS-CoV-2 infection, thus providing a bridge between studies with cell lines and studies in animal models.

## Materials and Methods

### Ethics statement

The study was performed in accordance with the Declaration of Helsinki, under a protocol approved by the University of Florida Institutional Review Board (IRB202000920) after written informed consent from participants.

### Recruitment, sample collection, and processing

Lung tissues were obtained from participants undergoing lung resection surgery or from allografts that underwent volume reduction before transplantation. In cases of lung resection for nodules, samples far from the site of pathology were utilized. Pleural tissue and staple lines were dissected away, and the lung tissue was then sectioned into approximately 0.5g pieces. The tissue was then minced with surgical scissors into samples with a mean diameter of 0.91 mm (range 0.40-1.5 mm), and equally distributed into a 24-well plate. Tissues in each well were resuspended in 1.2 mL of a commercial DMSO-free cryopreservation medium (CryoSOfree™, C9249; Sigma-Aldrich), divided into 200 μL aliquots, and then transferred to cryotubes each containing 800 μL of additional cryopreservation media. Samples were frozen at −80°C overnight, then transferred to vapor-phase liquid nitrogen storage.

### Viral cultures

HCoV-OC43 (strain HCoV-OC43-JAL-1) and SARS-CoV-2 (strain UF-1) viruses were propagated in Vero-E6 cell line (ATCC® CRL-1586™) in advanced DMEM (Gibco™, 12491015; Thermo Fisher) supplemented with 10% HyClone™ Defined low antibody, heat-inactivated, gamma-irradiated fetal bovine serum (SH30070.03IR2540; Cytiva), 1% L-alanine and L-glutamine supplement (GlutaMAX™, 35050061; Gibco™), and 1% penicillin/streptomycin mixture (17-602E; Lonza™ BioWhittaker™), at 37°C in 5% CO_2_ in 75 cm^2^ flasks. All SARS-CoV-2 work was performed in a BSL3 facility using approved protocols.

To inoculate the virus, media was aspirated from flasks with >80% confluent cell monolayers and replaced with 3 mL of fresh media. 1 mL of viral stock (containing 106 PFU) was added to each flask, and the flask was incubated at 37°C (33°C for hCoV-OC43) in 5% CO2 for 1 h, with manual rocking every 15 min. A “mock-infected” negative control cell culture was inoculated with 4 mL of media without virus and handled in parallel. The flasks were observed daily for development of virus-induced cytopathogenic effects (CPE) and harvested after 48–96 h of incubation, at which point the cells displayed ≥95% CPE and about 25% of the cells had detached from the growing surface.

H-CoV-OC43 viral stocks: The flasks were frozen once at −80°C, thawed to room temperature, scraped to lift off cellular debris and contents centrifuged at 2000 xg for 10 min at 4°C in a centrifuge bucket sealed with a safety cap. A sterile filtered solution of 60% w/v D(+)-Trehalose dihydrate (AC182550250; ACROS Organics™) dissolved in aDMEM was added to the supernatant to a final concentration of 10% w/v. Viral stocks were stored at −80°C overnight in 1 mL aliquots, and then transferred to vapor-phase liquid nitrogen storage. The infectious virus titer of each batch was determined via plaque assay.

SARS-CoV-2 viral stocks: Vero cells were inoculated in a BSL3 lab as described above and incubated with SARS-CoV-2 for 72 hours, at which point the supernatant was removed from the cell culture flask and transferred into a 50 mL tube. The tube was centrifuged at 2000g for 10 min at room temperature in a swinging-bucket centrifuge with a safety cap. The clarified supernatant was then frozen in 1 mL aliquots in externally threaded screw-cap tubes. All transfers of tubes in and out of the centrifuge bucket were done inside a BSC in a BSL3 facility. Virus titer was determined by plaque assay.

### Tissue culture, infection, and drug treatments

Lung samples: Tissue samples were thawed for 5 min at 37°C then transferred to a 15 mL tube containing 10 mL PBS using 1000 μL wide-bore pipette tips, centrifuged at 200 xg for 1 min, supernatant aspirated, and samples were resuspended in 1 mL PneumaCult media (PneumaCult-Ex, 05008; Stemcell Technologies) with 1% penicillin/streptomycin mixture (17-602E; Lonza™ BioWhittaker™), 50 ng/ml gentamycin sulfate (345815; Sigma-Aldrich) and 1.25 μg/ml ertapenem sodium (SML1238; Sigma-Aldrich). The concentration of tissue samples was measured by counting the number in 20 μL on a microscope slide, and 20-40 samples in 100 μL were added to individual wells of a 96 well plate. For studies of lung microtissues, ultra-low cell-binding flat-bottom 96-well cell culture microplates containing hydrogel (3474; Corning™) were hydrated with PBS for 30 min at 37°C. PBS was aspirated prior to addition of lung microtissues. One vial of cryopreserved lung tissue provided enough samples for approximately 10 wells at this concentration.

Infection and treatment of lung tissue: Lung samples were prepared in a BSL2 facility. In cases where drugs were added to the media, these drugs were added prior to the addition of virus. Prepared plates were transported to the BSL3 lab in a sealed secondary container inside a foam cooler. In the BSL3 lab, virus stocks were thawed and diluted to the desired concentration (ranging from 10^2^-10^4^ PFU/mL) in complete PneumaCult media in a BSC. Tissues were infected in a total of 150 uL of media for 24 hours. The inoculated plate was incubated at 37°C (33°C for hCoV-OC43 infections) in 5% CO2. In cases where the culture was longer than 24 hours, 120 uL of media was aspirated from each well and deposited in bleach to sterilize any infectious agents. An additional 120 uL of media containing the appropriate concentration of specified drugs was added to each well and cultured for an additional 24 hours. At the end of the experiment media was aspirated from each well and samples were washed 3X in PBS by adding 150 uL to each well and then removing as much of the liquid as possible prior to placing samples in 500 uL Trizol LS Reagent (10296-028; Thermo Fisher). A maximum of 100 uL of liquid + tissue samples was placed in each tube. For experiments taking place in the BSL3 lab, Trizol was transported in externally-threaded screw-cap tubes and disinfected prior to removal from the BSC. All samples were transported out of the BSL3 lab in disinfected sealed secondary containers containing absorbent material.

To study the effect of drugs on viral replication, the following drugs were used: β-d-N4-hydroxycytidine (9002958; Cayman Chemical), chloroquine (C6628; Sigma), water-soluble dexamethasone (D2915; Sigma), metformin (A10573; Adooq Bioscience), remdesivir (329511; Medkoo), and sirolimus (A10782; Adooq Bioscience). To calculate percent inhibition for remdesivir and sirolimus, which are soluble in DMSO, samples infected and treated with an equal volume of DMSO in the absence of drug to determine the maximum signal. All other drugs were water soluble and were compared to the infected but untreated control.

Vero-E6 cells used to test inhibition of viral growth were cultured to an estimated 90% confluence in 96 well plates prior to testing and then infected with 10^4^ PFU (approximating an MOI of 1) in viral growth media containing the specified drugs as described above. 10^4^ PFU was used for infection to enable a direct comparison between Vero cell cultures and lung microtissues infected with SARS-CoV-2. Media containing virus was removed after 24 hours and replaced with fresh media containing the specified drugs. After an additional 24 hours of culture, the media was aspirated and 100 uL of Trizol LS was added to each well to lyse the cells and capture the RNA. Trizol was then transferred into externally threaded screw-cap tubes containing an addition 400 uL of Trizol LS. Tubes were

### Flow Cytometry

Flow cytometry was only performed on uninfected cells. Biopsies were resuspended in RPMI-1640 (12-167F; Lonza™ BioWhittaker™) with 125 ng/mL Liberase (LIBTM-RO, 5401119001; Sigma-Aldrich) and 50 units/mL DNase I (D5025; Sigma-Aldrich), agitated for 1 h at 37°C, then titurated approximately 30x using a 1 mL syringe through an 18G needle to further disintegrate tissues forming a single-cell suspension. Cells were then centrifuged at 400 xg for 5 min at room temperature, washed twice in RPMI-1640, resuspended in flow cytometry buffer (1 ml PBS with 2% FBS and 1 mM EDTA), and filtered through 100 μm Nitex nylon mesh (57-103; Genesee Scientific). Concentration and viability were determined using a hemocytometer and Trypan Blue exclusion (1691049; MP Biomedicals).

The lung cells were washed and resuspended in 100 μL PBS and stained with a fixable viability dye (Zombie Aqua™, 423101; BioLegend) for 10 min at room temperature protected from light. Cells were washed, resuspended in 100 μL buffer, and then stained with antibodies against various surface cell markers (Table 1). During staining, serum from humans with AB blood group (HP1022; Valley Biomedical) was added to the samples to block non-specific binding. Surface marker antibodies were added at a concentration of 0.5 μL per 100 μL and then the samples were incubated for 20 min in the dark at room temperature on the orbital shaker. After staining, samples were washed 2x in flow cytometry buffer, centrifuged, and then fixed for 10 min in normal buffered formalin. The samples were again centrifuged at 400 xg for 5 min. The formalin was then aspirated, and the samples were washed 2x with PBS. Resuspended samples were analyzed on a BD Symphony instrument (BD Biosciences, San Jose, CA). Unstained cell samples from each donor and compensation beads from the Invitrogen™ AbC™ Total Antibody Compensation Bead Kit (A10497; Thermo Fisher) were used to set voltages and create single stain controls. Flow cytometry data was then analyzed using FlowJo X (BD biosciences).

**Table 1:** Antibodies and reagents used.

| Antibody | Clone | Supplier | Catalog Number |
| --- | --- | --- | --- |
| *APC anti-HLA-DR* | Immu-357 | Beckman Coulter | IM3635 |
| *APC/Cyanine7 anti-human CD206 (MMR)* | 15-2 | BioLegend | 321120 |
| *Brilliant Violet 711™ anti-human CD31* | WM59 | BioLegend | 303136 |
| *E-Cadherin* | 4A2C7 | ThermoFisher | 180223 |
| *FITC anti-human Podoplanin* | NC-08 | BioLegend | 337025 |
| *HCoV-OC43 Nucleocapsid protein* | 542-7D | Sigma | MAB9013 |
| *Pacific Blue™ anti-human CD11c* | 3.9 | BioLegend | 301626 |
| *PE anti-human CD45* | J33 | Beckman Coulter | IM2078U |
| *PE/Cyanine7 anti-human CD326 (EpCAM)* | 9C4 | BioLegend | 324222 |
| *SARS Nucleocapsid Protein (unconjugated)* | Polyclonal | NOVUS | NB100-56683 |
| *SARS Spike protein (unconjugated)* | Polyclonal | NOVUS | NB100-56578 |
| *Zombie Aqua™ Fixable Viability Kit* | -- | BioLegend | 423101 |
| *ZO-1 (unconjugated)* | Polyclonal | ThermoFisher | 40-2200 |

### Quantification of virus titers and cytokine responses

RNA was isolated from washed tissue samples using Zymo Direct-zol RNA MicroPrep kits according to the manufacturer’s instructions (11-330MB; Genesee Scientific). RNA concentration was quantified using a Nanodrop spectrophotometer (ThermoFisher), and 30 ng of RNA from each sample was reverse transcribed using the iScript™ cDNA Synthesis Kit (1708891; Bio-Rad). Real-time PCR was performed using Applied Biosystems™ TaqMan™ Gene Expression Master Mix (4369016; Thermo Fisher) with pre-developed primer/probe assays from ThermoFisher (CXCL8 Hs00174103_m1, IFNb1 Hs01077958_s1, IL6 Hs00174131_m1). Delta-delta CQ was calculated using the 18S ribosomal RNA primer/probe set (4331182; Thermo Fisher). Virus RNA (vRNA) levels corresponding to those encoding nucleocapsid proteins 1 and 2 were assessed using primer/probe mixes from IDT (2019-nCoV RUO Kit, 10006713) and a standard curve was prepared using the 2019-nCoV_N_Positive Control (10006625; Integrated DNA Technologies). For DMSO-soluble drugs (remdesivir and sirolimus) the inhibition of viral growth was calculated using infected samples treated with an equivalent amount of DMSO.

### Immunofluorescence staining

Samples were fixed in neutral buffered formalin (HT501128; Sigma-Aldrich) overnight at 4°C in an externally-threaded screw-cap tube prior to removal from the BSL3. Samples were then removed from the BSL3, and stored in PBS for 12-16 h at 4°C. Prior to staining samples were washed 2x in PBS, and allowed to equilibrate to room temperature for 1h. Samples were then permeabilized in PBS + 0.5% Triton X-100 (BP151-500; Fisher BioReagents) for 2 h, washed 2x in PBS, and blocked with 3.0% bovine serum albumin (A8806-5G; Sigma-Aldrich) in PBS for 3 h at room temperature. Samples were then incubated overnight with conjugated antibodies in sealed tubes at 4°C. The antibodies used in this study include E-cadherin, polyclonal nuclear capsid and spike SARS-CoV-2 antibodies, and HCoV-OC43 antibodies (see table 1). When staining for ZO-1, samples were incubated with primary rabbit anti-ZO-1 antibody overnight at 4°C, followed by washing and incubating with conjugated secondary antibodies (A-21433; Invitrogen) against the appropriate species for 3h at room temperature. Antibodies to viral proteins were directly conjugated using antibody labeling kits (ThermoFisher). For viability staining, a live/dead kit (R37601; Invitrogen) with Calcein AM and BOBO-3 Iodide was utilized following manufacturer protocol, and Hoechst 33342 (H3570; Invitrogen) was added to visualize nuclei. Lung tissue samples were then imaged using a Nikon A1R HD25 confocal microscope with high-definition galvano scanner.

### Statistical analysis

Data were analyzed using Prism software (version 9.0 for Mac, GraphPad, San Diego, CA, USA). Descriptive data were summarized as median and interquartile range (IQR). Two sample groups were compared using the Wilcoxon rank-sum test. Comparisons of multiple groups over a range of virus inocula was performed using two-way ANOVA with Sidak multiple comparison test when all groups were of equal size, or mixed-effects analysis when groups differed in size. In multiple comparison tests, multiplicity-adjusted *p* values (Dunnett test) are reported. Linear correlations between variables were assessed using Pearson coefficient. Probability value of <0.05 was considered statistically significant.

## Results

We recruited 16 subjects, most of whom had at least one risk factor for severe COVID-19 infection, including old age and comorbidities (Table 2).

**Table 2:** Characteristics of the clinical cohort used in Fig 3.

|  |  |
| --- | --- |
| Subjects, n | 16 |
| Median age (IQR) | 71 (64-76) |
| Male sex, n (%) | 11 (69) |
| Blood group, n (%) |  |
| O | 7 (44) |
| A | 6 (38) |
| B | 3 (19) |
| Smoking history |  |
| Never, n (%) | 3 (19) |
| Past, n (%) | 10 (63) |
| Active, n (%) | 3 (19) |
| Median pack-years in ever-smokers (IQR) | 20 (14-40) |
| Co-morbidity, n (%) |  |
| Active cancer | 12 (75) |
| Hypertension | 10 (63) |
| Over-weight or obese | 6 (38) |
| COPD | 2 (13) |
| Type II diabetes | 1 (6) |
| Congestive heart failure | 1 (6) |

COPD, chronic obstructive pulmonary disease; IQR, interquartile range.

### Composition of cryopreserved lung tissue

We began by assessing the viability and integrity of cryopreserved lung tissue after 48 hours of culture. We found a comparable viability to fresh tissue when cultured for 48 h as measured by flow cytometry (Fig 1A-B). All major subsets of cells were present in cryopreserved tissue, including epithelial cells, endothelial cells and leukocytes (Figure 1C-D). T cells, defined by CD3+ staining, comprised 6.3% of the total cell population, whereas alveolar macrophages composed 1.8% and other monocytic phagocytes made up approximately 0.3%. Each leukocyte population is consistent with previous results in which a collagenase-based method is used to digest lung tissue (10), but flow cytometric analysis showed type 1 epithelial cells to constitute an estimated 4% of total viable cells within the samples, lower than approximately 8% of lung tissue reported in literature (11, 12). We attribute this to the loss of fragile cells during processing for flow cytometry (13). To obtain a more accurate measurement of epithelial cell viability, we imaged the samples using confocal microscopy after staining with Calcein AM, a cell-permeable viability dye that only fluoresces in metabolically active cells, and found an abundant population of viable cells with type I epithelial morphology within the samples (Figure 1E). We confirmed this finding by staining the cryopreserved samples with zona occludens-1 (ZO-1), which identifies intact tight junctions between lung epithelial cells (14), and the epithelial cell marker E-cadherin (Figure 1F-G).

**Figure 1:**
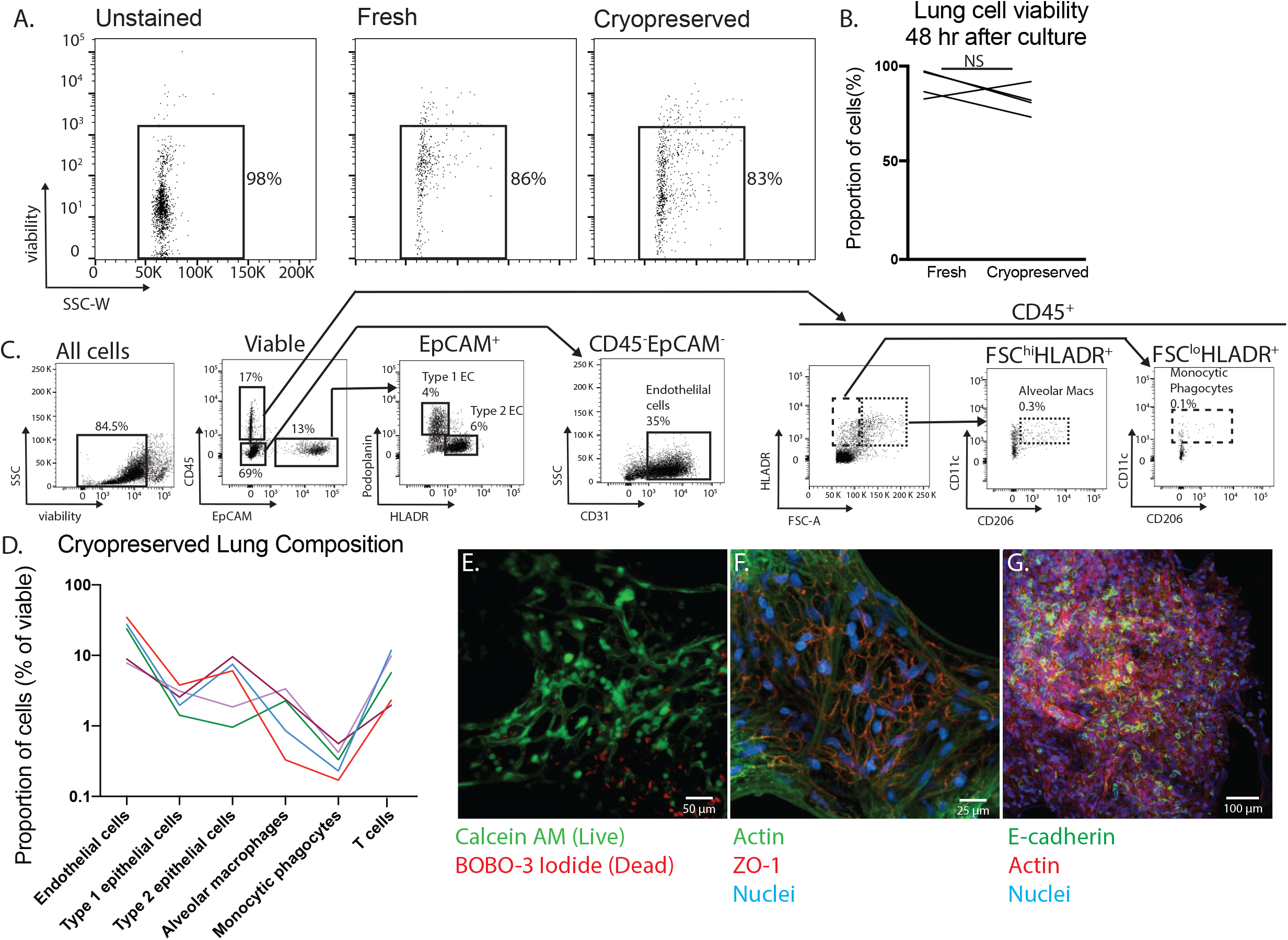
Lung microtissues are viable after cryopreservation. A) Flow cytometry was used to assess viability in microtissues cultured for 48 hours. Both fresh and cryopreserved tissues from the same donor were assessed using a fixable viability stain. B) Graph representing four matched samples. Each sample consists of between 20-40 microtissues cultured for 48 hours. *p*=0.25 as determined by Wilcoxon rank-sum test. C) The composition of cryopreserved lung was assessed by flow cytometry. Measured populations include epithelial cells, endothelial cells and monocyte/macrophage populations. D) The cellular composition of cryopreserved samples from five different donors was assessed using the gating strategy depicted in C. Each line represents the average population present in three samples, consisting of between 20-40 microtissues from each donor. E) Viability of cryopreserved tissues was assessed by microscopy using Calcein AM and BOBO-3 Iodide in microtissues cultured for 48 hours. F) Tight junctions in cultured microtissues were also assessed using Zo-1 with co-staining for both actin and nuclei using phalloidin and Hoechst 33342 respectively. G) Microtissues stained with E-cadherin demonstrate the presence of epithelial cell populations within the cryopreserved samples.

### In vitro infection of lung tissue with coronaviruses

We began by infecting lung samples from two donors with each of two human coronaviruses: HCoV-OC43, an endemic cause of the common cold (15, 16) that infects cells via cell membrane sialyl acid groups (17, 18), and SARS-CoV-2. Both HCoV-OC43 and SARS-CoV-2 could infect cryopreserved tissues in a dose-dependent manner (Figure 2A-B) and both viruses were detectable within the tissues by immunofluorescence microscopy (Figure 2C). Interestingly, SARS-CoV-2 protein was also detectable outside the tissue, consistent with viral shedding. Surprisingly, we found a marked disparity in the IL6 transcriptional response to both coronaviruses between the two donors, with a 100 fold increase in one and no change in the other (Figure 2D).

**Figure 2:**
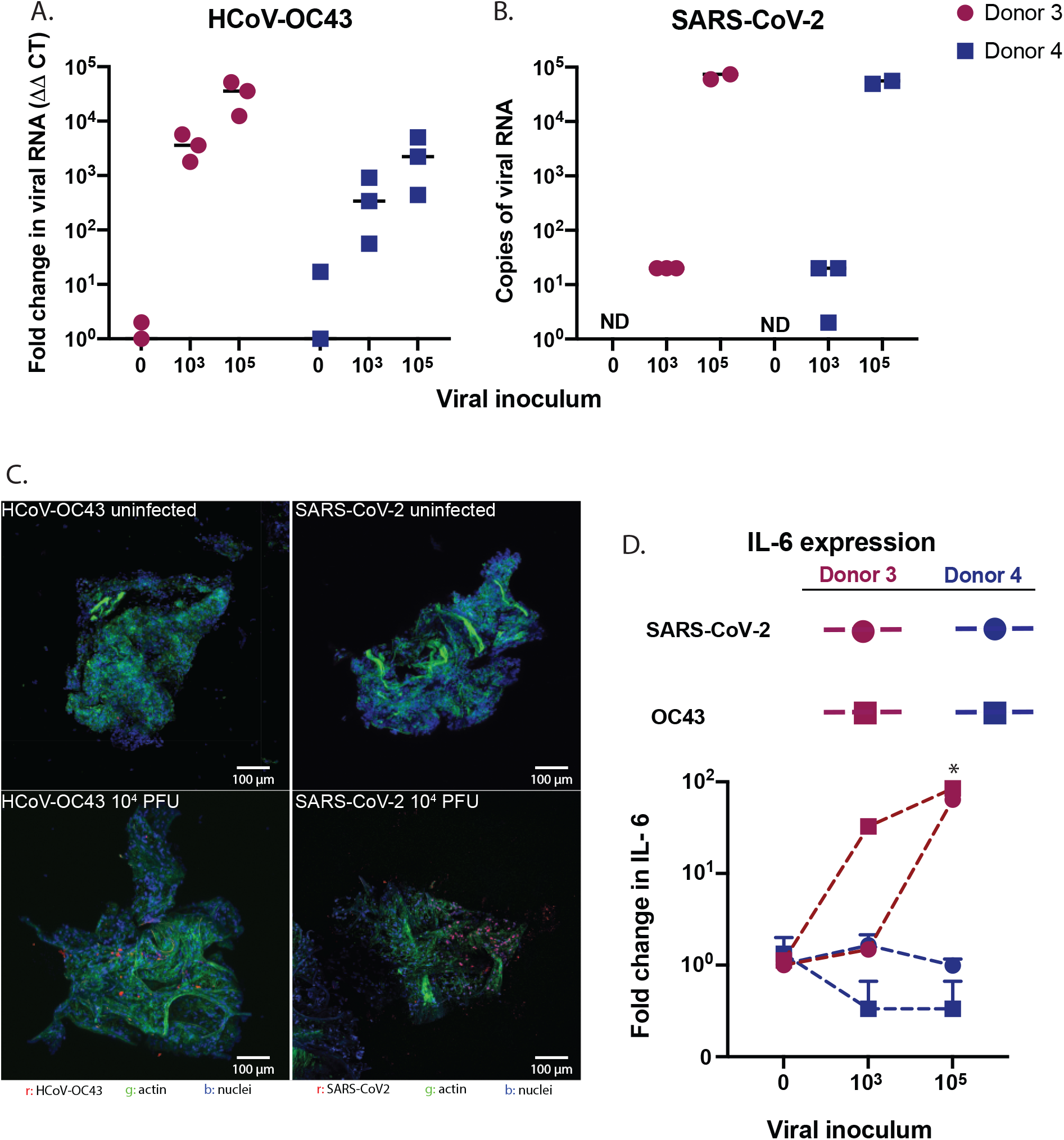
Lung microtissues can be infected with coronaviruses. A-B) Lung microtissues were infected with the specified PFU of either HCoV-OC43 or SARS-CoV-2 for 24 hours. The presence of virus in biopsies was assessed by qPCR. Two-way ANOVA analysis indicates a significant dose effect for both HCoV-OC43 (*p*=0.02) and SARS-CoV-2 (*p*=0.0001). HCoV-OC43 infection varied significantly in the two donors used (*p*=0.04) C) Micrographs of lung tissue stained with antibodies to HCoV-OC43 or SARS-CoV-2 and actin. D) The expression of *IL6* in the same samples as used in A-B. Two-way ANOVA indicated *IL6* expression in donor 3 was significantly increased at the 1×10^4^ dose (*p*=0.02).

We next sought to better assess this variance in the host response to SARS-CoV-2 infection within our tissue bank. As expected, we found an inoculum-dependent increase in viral protein transcription (Figure 3A-B). We selected expression of *IFNB1, IL6*, and *CXCL8* as markers of host response, representative of overlapping transcriptional responses to viral infection. We found highly heterogeneous responses among donors in response to the same viral inoculum, with a range of 0 to 34-fold induction in *CXCL8*, 0 to 85-fold induction in *IL6*, and 0 to 214-fold induction in *IFNB1* (Figure 3C-E). We found no correlation between the host response and age, sex, blood type, or presence of comorbidities in this cohort (data not shown). Comparison of cytokine induction within each donor showed that most donors displayed greater induction of cytokine transcription with higher viral inocula (Figure 3F). Within each donor, there was a strong correlation between the induction of *IL6* and *CXCL8* in response to both low and high viral inocula (*r^2^* >0.92), albeit with marked inter-individual variability, but no correlation between the expression of *IFNB1* with either *IL6* or *CXCL8* (*r^2^* <0.1; Figure 3F and supplemental figure).

**Figure 3:**
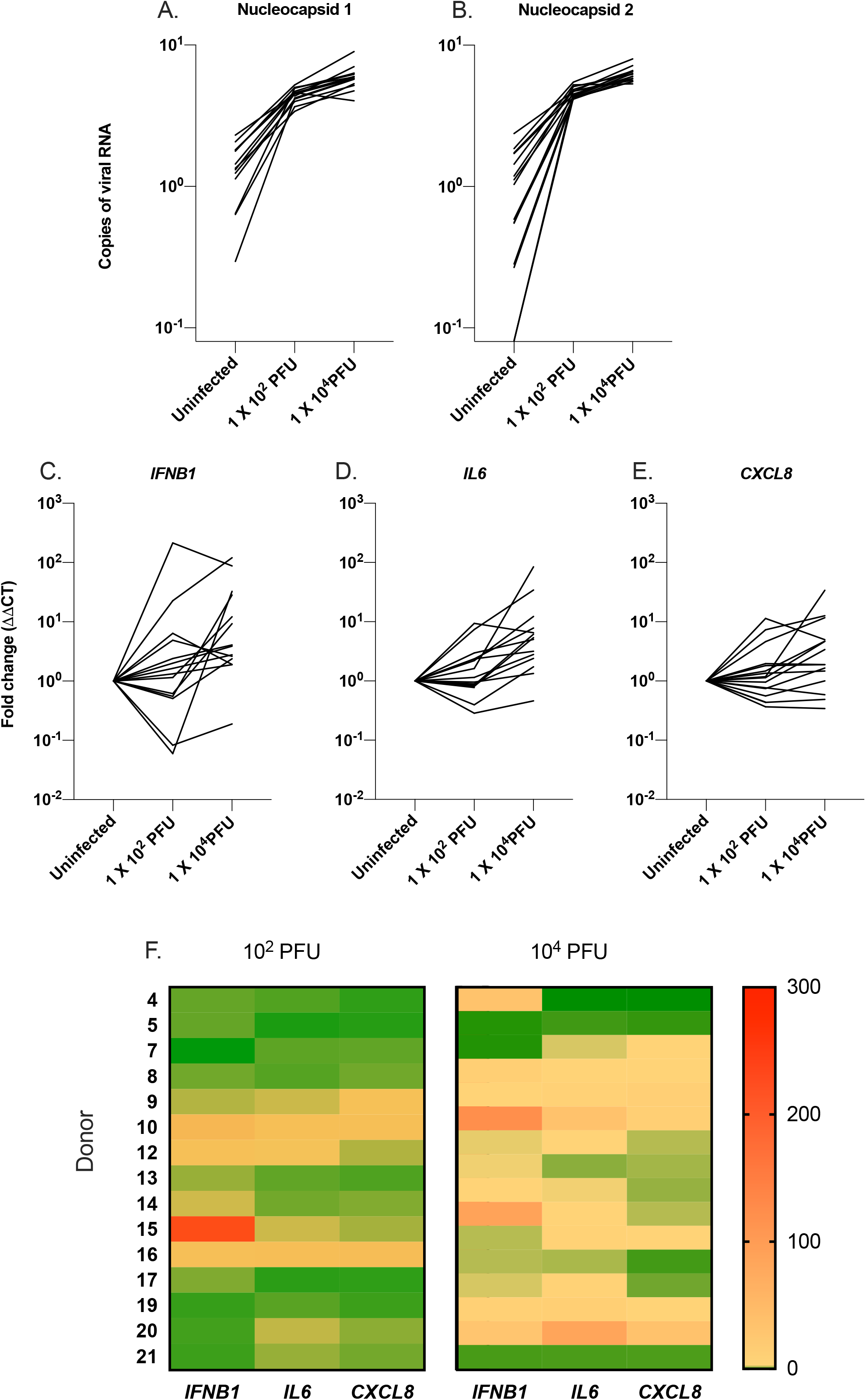
Heterogeneity in the host response to SARS-CoV-2 infection. A-B) Assessment of viral copy numbers in microtissues from different donors infected with SARS-CoV-2 for 24 hours. C-E) Measurement of the host response to infection, including expression of *IFNB1, IL6* and *CXCL8* in these same samples. In figures A-E, Two-way ANOVA indicates significant interaction between the donor and the virus inoculum (*p*<0.0001), with both factors contributing to the variance in the response. F) Heat maps comparing the level of cytokines in each donor at each dose of virus.

### Antiviral therapy in lung tissue culture

We reasoned that drug testing for COVID-19 relies on *in vitro* infection of cell lines, which may not accurately represent the response of human tissue to infection. To assess this, we next tested the effect of six drugs on SARS-CoV-2 viral titer in lung tissue from five different donors and VeroE6 cells. We found that dexamethasone, a drug found to improve outcomes in COVID-19 infection (19), reduced viral titers in all donor tissues, but not in VeroE6 cells (Figure 4A). In contrast, chloroquine reduced viral titer in VeroE6 cells but not in the lung tissue donors (Figure 4B), consistent with prior research demonstrating that chloroquine does not inhibit viral titer in lung epithelial cells (20, 21). Remdesivir, which has been variably effective in COVID-19 in clinical trials (22–24), did not significantly affect viral titer in any of the *in vitro* infections (Figure 4C).

**Figure 4:**
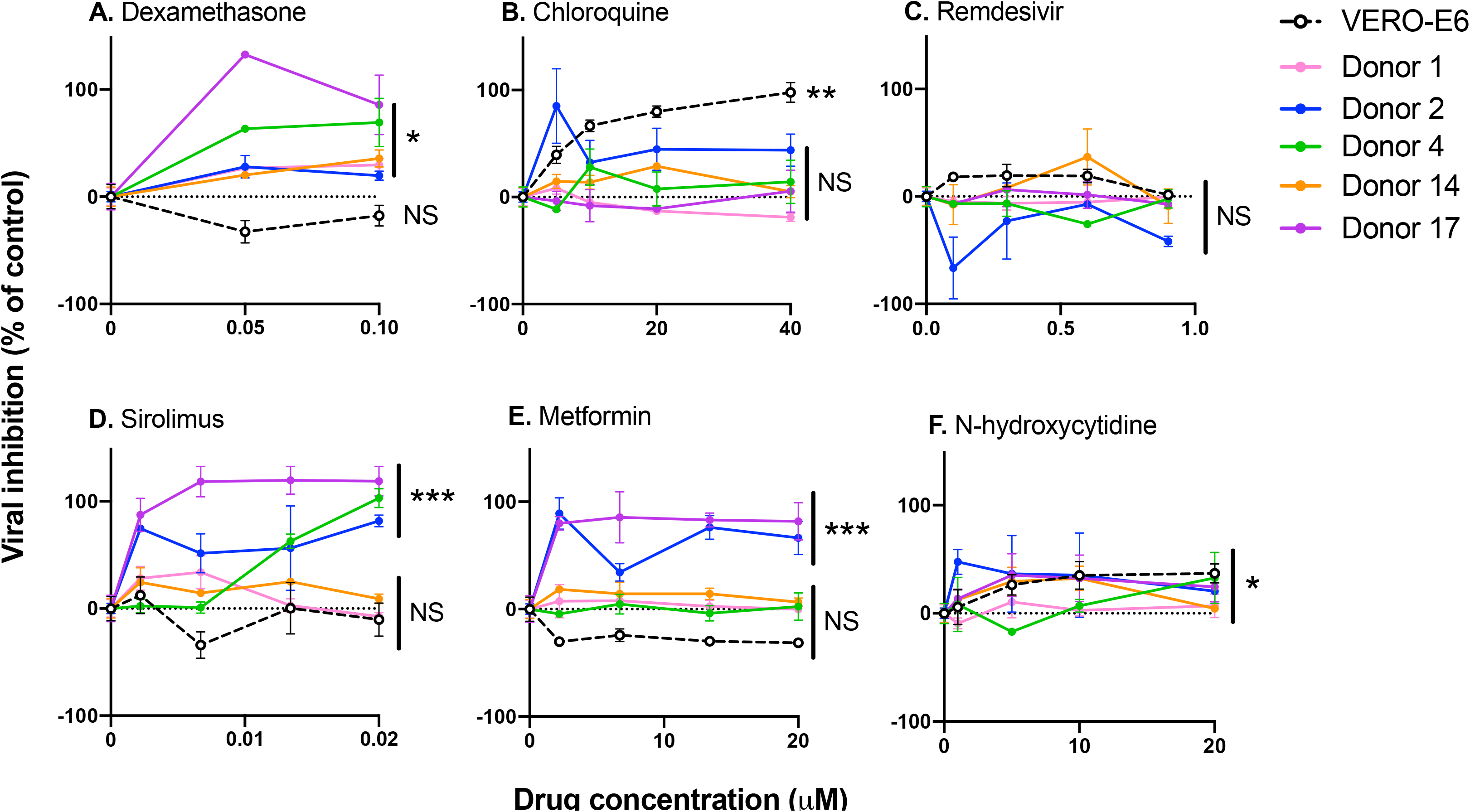
Effect of drug treatment on viral titer in lung microtissues. A-F) The percent inhibition in viral titer for six different drugs in five different donors and the Vero-E6 cell line. Significance was determined using a two-way ANOVA to test for significant interaction between individual donors and the dose of drug. \**p*<0.05; \*\**p*<0.01; \*\*\**p*<0.0001.

Finally, we tested three investigational therapies previously predicted as potentially effective in SARS-CoV-2 infection, including sirolimus (25, 26), metformin (27–29), and N-hydroxycytidine (30). We found that sirolimus was highly effective in reducing viral titer in tissues from three donors, but did not significantly affect titer in two donors or Vero cells (Figure 4D). A similar heterogeneity was observed with metformin (Figure 4E). N-hydroxycytidine, an experimental drug that inhibits viral transcription (30), resulted in a significant reduction in viral titer in all tested samples (Figure 4F). These results highlight the considerable heterogeneity of antiviral host response and medication effectiveness in human tissues.

## Discussion

The *in vitro* study of human lung tissues has been hampered by difficulties in growing and maintaining 3-Dimensional tissue in culture. The current technologies to achieve this, namely lung-on-a-chip, lung organoids, and precision-cut lung slices, have provided invaluable insights to our understanding of the human lung. While each of these technologies has its strengths and limitations, all are limited by the need for fresh tissues. Our study demonstrates that polyampholyte-based cryopreservation media can be used to preserve normal and diseased lung tissues in large batches. After cryopreservation, thawed microtissues displayed excellent viability, were metabolically active, and contained the major cell populations of the human lung. Our data is in agreement with previous studies demonstrating the advantages of using polyampholyte-based media as a cryopreservative (8, 9).

A notable finding in our study was that the human tissues tested displayed similar infectability after inoculation with a given SARS-CoV-2 dose, as measured by expression of viral nucleocapsid proteins, suggesting that inter-individual variability in infection is not attributable to the susceptibility of host respiratory tissues to infection (Figures 3A-B). In contrast, infection with a given virus inoculum resulted in dramatic inter-individual heterogeneity in the cytokine responses of the infected tissues (Figures 3C-E). In this context, multiple studies have described the variability in host susceptibility to COVID-19 infection, using clinical outcomes, cytokine responses, and duration and extent of viral shedding as readouts, leading to the identification of acquired polymorphisms as risk factors for severe disease (31–37). Our data adds to this literature by assessing the effect of a uniform viral inoculum between hosts, allowing us to distinguish between tissue susceptibility to infection as opposed to antiviral and inflammatory host responses.

Another important component of our work was the demonstration of antiviral drug testing using primary human lung tissues, which represents an advance over the study of drugs in the context of non-physiologic cell lines. In this context, we confirmed the *in vivo* efficacy of dexamethasone and the ineffectiveness of chloroquine in suppressing viral growth in human tissues. An unexpected finding in our work was the lack of effect of remdesivir in both VeroE6 cells and human tissues, in contrast to prior studies in several cell lines (38–40). One potential explanation for this discrepancy is that these studies used a much lower MOI (of 0.05 to 0.2), whereas we used an MOI of one in Vero cell cultures in order to inoculate with a virus titer equal to that used in our lung culture system. In this context, one study demonstrated that remdesivir exhibited potent antiviral activity in primary airway epithelial cells cultured using an air-liquid interface but was not effective in reducing viral titer in Vero cells, similar to our findings (38). Overall, the difference in experimental conditions likely explain the variance in the published literature regarding the effectiveness of remdesivir *in vitro*.

Our *in vitro* lung tissue assay allowed us to test the predictions of a biophysical model for the SARS-CoV-2 life cycle (41). This model predicted that targeting of transcription and translation are especially sensitive processes that could allow effective antiviral targeting of SARS-CoV-2. Consistent with the model predictions the transcription inhibitor N-hydroxycytidine and the translation inhibitors metformin (41,42) and sirolimus were effective in inhibiting viral replication for at least some of the patients. Also consistent with the model predictions, the viral entry inhibitor, chloroquine, was not effective. The model failed to predict the lack of effectiveness of remdesivir, which may be due to the specific properties of the drug binding in the cells (41). The model did not make any specific prediction about dexamethasone, which targets immunomodulators. Overall, the biophysical model points to the potential effectiveness of targeting protein synthesis, motivating clinical testing of sirolimus and metformin, in particular.

We recognize a number of limitations in our study. First, lung tissue is currently only viable for up to 96 hours in our culture system, thereby limiting the observations regarding pathogenesis of infection to this short time frame. Second, like all *in vitro* systems, this culture system includes non-physiologic features that may impact the biology of infection, such as non-physiologic culture media and the absence of air. Third, the system does not capture the biology of recruited leukocytes, which undoubtedly play an important role in the development of lung injury in COVID-19. Fourth, the number of tissue donors in our study was limited to only 16, and serves only as a proof-of-principle. Our study was limited in its ability to detect host factors that may predispose to tissue infectability, aberrant host response, or effectiveness of antiviral drugs.

The current work suggests a number of avenues for further research. First, the ability to infect human tissues with SARS-CoV-2 *in vitro* allows for a systematic study of early events in different cell types in the lung, and comparisons between different hosts, potentially leading to new insights into disease pathogenesis and heterogeneity of disease phenotype in patients. Second, the current system provides a platform for the study of the effectiveness of other antiviral drugs in human tissues, in effect prioritizing drugs for *in vivo* testing or clinical trials. Third, *in vitro* infection of human lung tissue allows for studies to define mechanisms of action for medications. For example, the mechanisms that lead to inhibition of viral titer in response to sirolimus, metformin, or dexamethasone are unknown. While several studies have proposed mechanisms for metformin and sirolimus (25–27), dexamethasone is thought only to inhibit the host response. Our data indicate that early viral replication in lung tissue may be linked to a corticosteroid-sensitive mechanism that involves the host response to infection.

In summary, we provide evidence that human lung tissue can be utilized *in vitro* for the short-term study of SARS-CoV-2 infection. A lung tissue bank may also serve as a tool for the study of lung biology and drug screening beyond COVID-19, and facilitates the screening of multiple donors in parallel for purposes of toxicity and efficacy in primary human tissues. We envision that screening therapeutic agents in primary human tissue will take place after screening in cell lines, and in parallel with testing in animal models.

---

**Figure S1:** Correlation in host response to SARS-CoV-2. A-E) The correlation between the production of IL-6, CXCL8 and IFN-beta in each donor was assessed by linear regression for two doses of virus. R squared values are depicted in the bottom of each graph.

